# Dimerization of rhomboid protease RHBDL2 in lipid membranes addressed by FRET with MC simulations

**DOI:** 10.1101/292318

**Authors:** J. Škerle, J. Humpolíčková, P. Rampírová, E. Poláchová, L. Adámková, A. Suchánková, D. Jakubec, K. Strisovsky

## Abstract

Many membrane proteins are thought to function as oligomers, but measuring membrane protein dimerization in native lipid membranes is particularly challenging. Förster resonance energy transfer (FRET) and fluorescence correlation spectroscopy (FCS) are non-invasive, optical methods of choice that have been applied to the analysis of dimerization of single-spanning membrane proteins. The effects inherent to such two-dimensional systems, such as excluded volume of polytopic transmembrane proteins, proximity FRET, and rotational diffusion of fluorophore dipoles, complicate interpretation of FRET data and have not been typically accounted for. Here, using FRET and FCS we introduce methods to measure surface protein density and to estimate kappa squared, and we use Monte Carlo simulations of the FRET data to account for the proximity FRET effect occurring in confined 2D environments. We then use FRET and FCS to analyze the dimerization of human rhomboid protease RHBDL2 in its native lipid membranes. While previous reports have proposed that rhomboid proteases dimerize and this allosterically activates them, we find no evidence for stable oligomers of RHBDL2 in lipid membranes of human cells. This indicates that the rhomboid transmembrane core may be intrinsically monomeric. Finally, our findings will find use in the application of FRET and FCS for the analysis of oligomerization of transmembrane proteins in lipid membranes.

## Introduction

Membrane proteins frequently form functionally important homo or hetero-oligomeric complexes, and their surrounding milieu, e.g. plasma membrane, can shape their properties and interactions profoundly (1). It is thus important to study membrane protein interactions directly in their native membranes in a minimally invasive way. Optical, fluorescence-based methods appear ideal for this purpose. Fluorescence resonance energy transfer (FRET) is a powerful technique for measuring distances at the nanometer scale and thus it can potentially address protein oligomerization. FRET reports on the vicinity of protein which carries an energy donor in the presence of an acceptor, typically provided by fluorescent protein reporters, fused genetically to the protein of interest. In the three dimensional space (in solution), FRET response is observed almost exclusively when the proteins of interest are indeed physically interacting in oligomers. However, in a two dimensional confinement, of membranes, protein density can be elevated to such a level that even if the two proteins do not physically interact in a complex, energy can be transferred to multiple acceptor molecules that are in close proximity of the donor (2). Studying protein oligomers within cellular lipid membranes by FRET is therefore ideal for situations of low protein density, i.e. the interaction has to be rather strong. Measuring dissociation constants of weaker interactions requires higher protein surface density, which results in increased FRET ‘background’ arising merely from the proximity of non-interacting fluorescence acceptors (3, 4). We aimed at analyzing and compensating these and other limitations to allow usage of fluorescence techniques for the study of oligomerization of polytopic membrane proteins spanning a wide range of dissociation constants in live cell derived membranes.

Already under mild overexpression conditions common in cell-biological experiments, the ‘proximity-induced FRET’ effect mentioned above may become significant, and it has to be properly taken into account to avoid spurious results. This problem is typically circumvented by the use of FRET controls, i.e. validated known interacting and non-interacting protein pairs. In addition to that, protein density level in the membrane has to be evaluated and only samples with similar surface densities of donor and acceptor labelled proteins should be mutually compared. In most membrane structures of living cells, direct protein surface density determination is not feasible, since the membrane area from which the fluorescence signal is collected cannot be measured. Therefore, fluorescence intensity of the donor and acceptor is used instead for relative comparison (5). Even such a comparison, however, requires careful interpretation because the level of FRET is influenced not only by oligomerization and density-induced proximity, but also by the length of the linker between the protein of interest and the fluorescent probe. In addition, it is affected by the excluded area of studied proteins (6). Importantly, global protein surface density at the membrane can be determined in spherical giant plasma membrane-derived vesicles (GPMVs) formed from cells expressing proteins of interest (7-10). Finally, recent work analyzing the FRET efficiency signatures of non-interacting membrane proteins both experimentally and by Monte Carlo (MC) simulations indicates how the overall FRET efficiency should be corrected for the ‘proximity-induced FRET’ effect mentioned above (2).

Here we focus on intramembrane proteases of the rhomboid family, proteins formed by six or seven α-helical transmembrane segments (reviewed in 11). These enzymes occur widely across evolution and regulate EGF receptor signaling, mitochondrial dynamics, mitophagy, apoptosis, or invasion of apicomplexan parasites, and are relevant for a growing number of diseases including malaria, cancer and Parkinson’s disease (reviewed in 12). Based largely on *in vitro* experiments in detergent-solubilized state of three bacterial rhomboid proteases, these enzymes have been proposed to form stable dimers and thereby become allosterically activated (13). This is a significant mechanistic feature impacting on the interpretation of biological mechanisms involving rhomboids. Rhomboid proteases are members of a larger superfamily of rhomboid-like proteins (reviewed in 14) including pseudoproteases Derlins and iRhoms of high biological and medical importance. All superfamily members share the basic transmembrane fold, and it is thus important to establish whether regulatory dimerization is a common property of the superfamily or whether it is peculiar only to some subgroups. Here we zoom in on the well-characterized human rhomboid protease RHBDL2 (15-20), which is localized to the plasma membrane and consists of seven transmembrane helices, and analyze its oligomeric status in living cells.

We show that at low protein expression that allows for a single molecule approach, no RHBDL2 dimers are observed by FCCS. At higher protein densities required to reveal possibly weaker interactions, the use of FRET is needed. To this end, we use human cells expressing fluorescently labelled RHBDL2 to create GPMVs, which have simple geometry and show homogeneous distribution of the observed molecules. We then employ FRET in individual vesicles in combination with Monte Carlo (MC) simulations to investigate the oligomeric state of RHBDL2. In doing so we build on the MC-FRET approach introduced by Johannsson *et al* (21, 22) and later applied to lipid clustering by others (23, 24). Within this context we develop and implement several novel considerations that are essential when addressing large membrane inclusions (such as RHBDL2) linked to fluorescence proteins. In particular, we pay attention to the orientation factor kappa squared and propose a simple experimental approach to estimate its value in a relevant model system. We propose a single-molecule based microscopy approach to calculate surface density of fluorescent proteins, and we use Monte Carlo simulations to model the FRET measurement in every GPMV, thus accounting for the lack of *à priori* knowledge of the dissociation constant and the geometry of the putative dimer. Comparing the simulation read-out with the measured data then allows drawing conclusions on the dimerization of the membrane protein. In summary, we address several key limitations of using FRET for oligomerization studies of polytopic membrane proteins in live cell derived membranes and demonstrate their use by showing the lack of dimerization of rhomboid protease RHBDL2, which we confirm independently by cell biological ‘relocalization’ experiments. Our results can find use in the application of FRET and FCCS for the analysis of oligomerization of transmembrane proteins in lipid membranes.

## Materials and Methods

### Materials

Oligonucleotides were from Sigma-Aldrich (USA), KRD (CZ) or Generi Biotech (CZ), and restriction endonucleases and other enzymes for DNA cloning were purchased from New England Biolabs (USA) and Thermo Fisher Scientific (USA). All other chemicals were from Sigma Aldrich unless stated otherwise.

### Cloning

The fluorescence reporters eGFP and mCherry were cloned as fusions to the N-terminus of human RHBDL2 (NCBI Reference Sequence NP_060291.2) in the pEGFP vector (17). For relocalization experiments, DNA sequence encoding the peptide KDEL preceded by a (GS)_3_ linker was cloned at the 3’ end of the RHBDL2 gene in the eGFP/mCherry-RHBDL2 constructs. The constructs encoding murine CD8α fused to eGFP or mCherry were generated by cloning DNA fragments encoding CD8α-GSGGGS-eGFP/mCherry between the *EcoRI* and *BamHI* restriction sites of eGFP-pXJ41 (25, 26). The constructs encoding His_6_-eGFP or His_6_-mCherry were kind gifts of Evzen Boura and were generated by cloning the eGFP or mCherry encoding fragments into the pHis_2_ vector downstream of a His_6_ purification tag and TEV cleavage site (27). All constructs were verified using Sanger sequencing.

### Protein expression and purification

His_6_-eGFP and His_6_-mCherry were expressed from their T7 driven vectors in *E.coli* BL21(DE3) at 20 °C, inducing by 0.5 mM IPTG for 12 hrs. Cells were broken by 3 passages through Emulsiflex C3 (Avestin, Canada) in the presence of serine protease inhibitor phenylmethylsulfonylfluoride (at 1 mM), the insoluble fraction was removed by centrifugation at 15000×g for 30 min at 4 °C. The His-tagged proteins were purified from the supernatant by metal-chelate affinity chromatography using NiNTA agarose (Qiagen), and eluted into 20 mM HEPES pH 8.0, 300 mM NaCl, and 10% (w/v) glycerol using 250 mM imidazole, which was immediately removed by desalting into PBS pH 7.4 using a PD-10 desalting column (GE Healthcare). Protein concentration was determined from absorbance at 280 nm, and purified proteins were flash-frozen into liquid nitrogen and stored at −80 °C.

### Cell culture

HeLa cells (ATCC) were cultured in DMEM medium (Invitrogen) supplemented with 10% foetal calf serum (Thermo) at 37 °C and 10% CO_2_. For transfection, 2×10^5^ cells were seeded in 4 chamber dish (Cellvis, cat. no. D35C4-20-1.5-N) and transfected by FuGene6 (Promega). The DNA amounts transfected per well were: 50 ng of eGFP-RHBDL2 and 50 or 500 ng of mCherry-RHBDL2 fusion construct plasmids (for donor:acceptor ratios 1:1 and 1:10, respectively) for FLIM-FRET experiments, and 50 ng of FP-RHBDL2 vector with corresponding 50 ng FP-RHBDL2-KDEL fusion construct plasmids for the relocalization experiments. The transfection mixtures were complemented with empty vector pcDNA 3.1 to 1 μg of total DNA mass.

Spontaneously immortalized human keratinocytes HaCaT (28) (item no. 300493, Cell Lines Service, Germany) were cultured in DMEM medium (Invitrogen). To deplete endogenous RHBDL2, HaCaT cells were transduced with a recombinant lentivirus expressing the shRNA #01 targeting RHBDL2 (16) as described (15) and selected for puromycin resistance.

### Immunoblotting

The integrity of the expressed fusion proteins and the expression levels of RHBDL2 were examined by immunoblotting. Cell lysates were separated by 4-20% gradient Tris-Glycine SDS PAGE (BioRad), electroblotted onto a PVDF membrane (Immobilon-FL, Millipore), optionally stained by the Revert protein stain (Li-Cor) and scanned using infrared fluorescence scanner Odyssey CLx (Li-Cor), destained, blocked in the Casein Blocker solution (Thermo) for 1 hr at 25°C, and exposed to primary antibodies over night at 4°C. Primary antibodies were from rabbit, and were used at the following concentrations: α-RHBDL2 (Proteintech, cat. no. 12467-1-AP) at 1:500, α-eGFP (Cell Signaling Technology, cat. no. 2956) at 1:1000, α-RFP (Thermo Fischer Scientific, cat. no. R10367) at 1:3000 and secondary antibody Donkey α-Rabbit IgG (H+L) Cross-Adsorbed conjugated to DyLight 800 (Thermo Fischer Scientific, cat. no. SA5-10044) at 1:10000. Secondary antibody fluorescence was visualized using near-infrared fluorescence scanner Odyssey CLx (Li-Cor), and optionally quantified using Image Studio™ Lite (Li-Cor).

### GPMV preparation

GPMVs were prepared as described elsewhere (29). Briefly, HeLa cells transiently expressing eGFP and mCherry fusions to human RHBDL2 were washed by the GPMV buffer (10 mM HEPES, 150 mM NaCl, 2 mM CaCl_2_, pH 7.4). Then 2 mM solution of N-ethylmaleimide in the GPMV buffer was added and the cells were incubated for minimum 1 hour at 37 °C. Once GPMVs were formed, they were immediately used for microscopy experiments *in situ*.

### GUV preparation

GUVs were prepared by electroformation (30). Chloroform-lipid mixture was prepared so that total lipid concentration was 5 µg/µL, containing 75 mol % of POPC, 20 mol % cholesterol and 5 mol % DGS-NTA(Ni) (all lipids purchased from Avanti Polar Lipids, Alabaster, AL). Ten µL of the mixture was spread on two ITO coated glass electrodes each. The electrodes were dried under vacuum overnight and then assembled in parallel into a homemade Teflon holder containing 5 mL of 600 mM sucrose solution. For the electroformation, 10 Hz, harmonically oscillating voltage of 1 V peak-value was applied to the electrodes for 1 hour in an incubator set to 60 °C. For the imaging, BSA-coated 4-chamber glass bottom dish (In Vitro Scientific) were used, and 100 µL of GUVs were mixed with 100 µL iso-osmotic buffer (25 mM Tris pH 8, 10 mM MgCl_2_, 20 mM imidazole, 261.5 mM NaCl, 2 mM β-mercaptoethanol) containing the His-tagged fluorescent proteins.

### Microscopy

All the microscopy images were acquired on the LSM 780 confocal microscope (Zeiss, Jena, Germany) using 40×/1.2 water objective. For the FLIM-FRET and FCCS measurements the external tau-SPAD detectors equipped with time-correlated single photon counting (TCSPC) electronics (Picoquant, Berlin, Germany). For FCCS experiments, eGFP was excited with the 490 nm line of the Intune laser (Zeiss) pulsing at 40 MHz repetition frequency and mCherry was excited continuously at 561 nm. The excitation light was focused on the apical membrane of giant liposomes. The precise positioning was checked by maximizing fluorescence intensity and the apparent molecular brightness (31). Fluorescence intensity, collected by the same objective lens, was re-focused on the pinhole (1 airy unit) and the re-collimated light behind the pinhole was split on the external tau-SPADs in front of which emission band pass filters 520/45 and 600/52 for eGFP and mCherry signal, respectively, were placed. The intensity of the excitation light at the back aperture of the objective was 2 and 6 µW for 490 nm and 561 nm laser line, respectively. The collected data were correlated by home-written script in Matlab (Mathworks, Natick, MA) according to a described algorithm (32, 33). To avoid detector crosstalk, the red channel fluorescence signal was split according to its TCSPC pattern (exponential for the signal generated by the pulsed Intune laser and flat for the 561 nm continuous wave laser) into two contributions and only the signal assigned to the flat TCSPC profile was correlated. The data were processed as described (33). The FLIM-FRET data were acquired in the equatorial plane of giant liposomes. The data were accumulated during fast repetitive frame scans. Frame-sequentially, the acceptor signal was collected. The mean fluorescence signal of the donor and acceptor was used to obtain information on their concentrations by extrapolation of the calibration measurement as described in the Results section. The excitation intensity of the 561 nm laser was decreased to 0.5 µW in order to minimize photobleaching of the acceptor.

### Monte Carlo (MC) simulations

For each FLIM measurement carried out on a single GPMV/GUV, an MC simulation was executed. The input parameters for the simulation were the surface concentrations of donor- and acceptor-labelled proteins that were obtained directly during FLIM acquisition with the help of the calibration measurement (see further). The simulations were run for the following varying parameters: i) Förster radius (*R*_0_), for *κ*^2^ determination, or ii) dissociation constant, *K*_D_, and excluded radius, *R*_excluded_, when oligomerization was addressed.

MC simulations were performed on the grid of 1000 × 1000 pixels. Size of the pixel was set to be 0.1×*R*_0_. First, the total amount of proteins in the simulated field was calculated. Based on the assumed *K*_D_, overall numbers of monomers and dimers were calculated. In our planar system, the dissociation constant *K*_D_ is defined as follows:

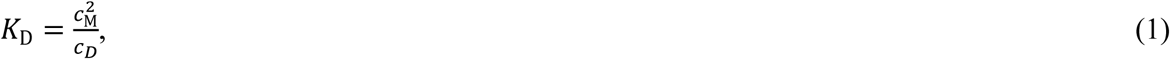

where *c*_D_ and *c*_M_ are surface concentrations of dimer and monomer, respectively.

All the monomers and one of the partners from each dimer were placed to the field so that they did not overlap (each occupies a circular area of a diameter of an excluded distance), i.e. a random position was generated and if it was occupied, it was rejected and another one was generated. Subsequently, the partners in the dimers were located: a random, already localized molecule was selected and a new one was placed directly next to it (so that the distance of their centers equals the excluded distance) at a random angle. Once the molecule was located, a random number was generated that distinguishes whether the molecule is a donor or an acceptor based on the experimentally known ratio between those. The system uses periodic boundary.

Once the molecules were localized in the field, a random donor was selected and overall FRET rate Ω_i_ was determined according to the formula:

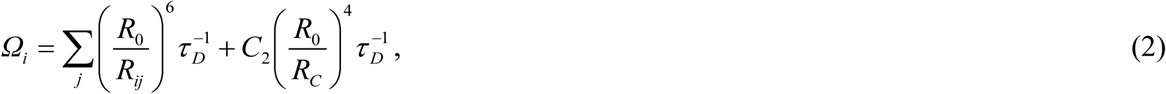

where *R*_ij_ is the distance between the selected donor and all acceptors closer than the cut-off distance *R*_C_ (10×*R*_0_), *τ*_D_ is the lifetime of the donor in the absence of acceptor, and *C*_2_ is the number of the acceptors inside the circle of the Förster radius (21, 22, 34). Then a random time Δ*t*_i_ at which the energy transfer occurs was generated: 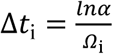, where *α* is a random number from the interval of 0 to 1. For each situation, i.e. each GPMV/GUV, 100 configurations of donors and acceptors were generated and at each configuration 100 random excitations were performed. From the final histogram of Δ*t*_i_, the donor decay in the presence of acceptors *I*_DA_(*t*) and the FRET efficiency η was generated:

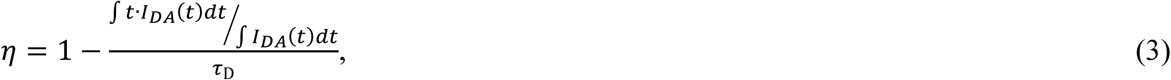

where *τ*_D_ is the fluorescence lifetime of donor in the absence of acceptor.

FRET efficiencies obtained from the simulations (η_D,A_) for the given input parameters (Förster radius *R*_0_, or *K*_D_ and excluded distance *R*_excluded_) were compared to the measured data (η_D,A exp_) in terms of χ2:

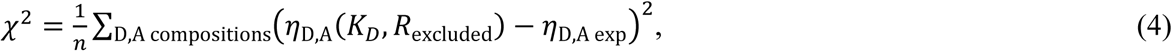

where *n* is the number of GPMVs/GUVs measured in the experiment, i.e. all experimental donor/acceptor compositions.

## Results

### Investigating dimerization of human rhomboid protease RHBDL2 by fluorescence cross-correlation spectroscopy

The joint motion of proteins of interest provides the most convincing evidence of their physical interaction, and single-molecule techniques based on tracking individual fluorescently labeled proteins thus represent straightforward tools to address protein-protein oligomerization. Since single-molecule based approaches require protein concentrations low enough to distinguish individual molecules, they are applicable only to strongly interacting species (i.e. those that form significant fraction of oligomers at low total concentrations).

We have first employed fluorescence cross-correlation spectroscopy (FCCS) (35) to investigate the dimerization status of human RHBDL2 in live cell derived membranes of GPMVs. The FCCS technique analyzes fluorescence intensity fluctuations arising from individual molecular transits through the diffraction limited laser focus, and the temporal cross-correlation between the signal of fluorescence reporter labeled membrane proteins suggests their joint motion. We have performed the experiment in the apical membrane of GPMVs (Fig. 1J) derived from HeLa cells co-expressing eGFP-RHBLD2 and mCherry-RHBDL2, both localized largely to the plasma membrane and mostly full-length (Fig. 1A, D, G). Surprisingly, the auto- and cross-correlation functions (Fig. 1M) did not suggest any interaction of RHBDL2 monomers at this concentration range. The same experiment was performed with murine CD8α (Fig. 1B, E, H, K, N), which is known to form dimers and localize to the plasma membrane (36). As expected, the co-expressed eGFP and mCherry fusions of CD8α exhibited a positive cross-correlation amplitude (Fig. 1N), indicative of a dimer. In contrast, eGFP and mCherry proteins attached to the membrane via Ni-NTA lipid containing GUVs by a His-tag (Fig. 1C, F, I, L) did not show any cross-correlation (Fig. 1O), serving as negative, monomeric controls. Taken together, these results indicated that if RHBDL2 dimers exist at all, their interaction is relatively weak, and their analysis thus requires methods that can access higher concentration ranges than FCCS measurements.

**Fig. 1:**
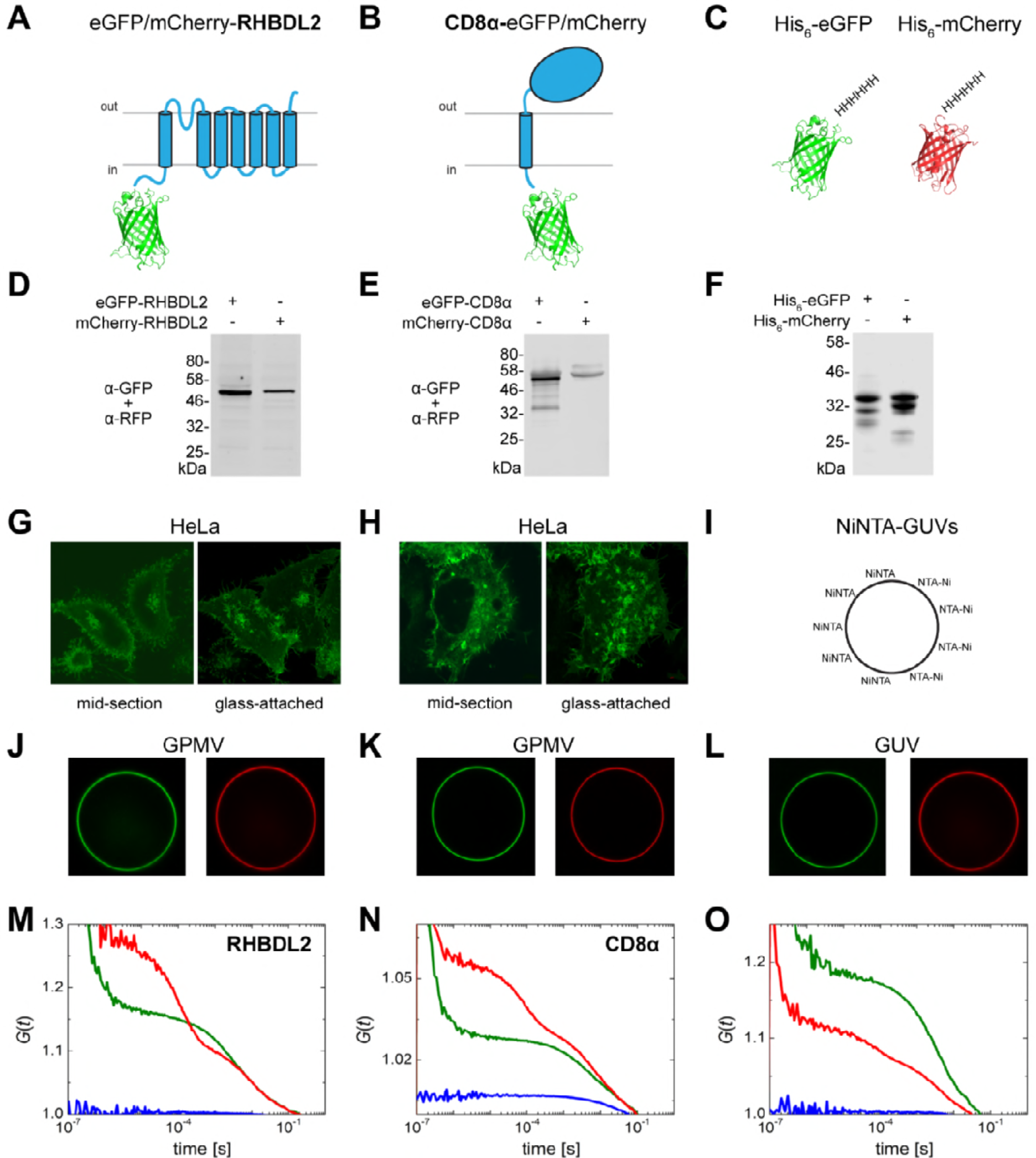
Fluorescence cross-correlation analysis of dimerization of RHBDL2 and CD8α. Construct scheme (**A**), Western blot showing expression and integrity (**D**), live-cell GFP fluorescence showing subcellular localization (**G**) and fluorescence images of GPMVs (**J**) of eGFP-RHBDL2 and mCherry-RHBDL2; **M**, Auto- and cross-correlation functions of eGFP-RHBDL2 and mCherry-RHBDL2 in GPMVs; Construct scheme (**C)**, Western blot showing expression and integrity (**E**), live-cell GFP fluorescence showing subcellular localization (**H**) and fluorescence images of GPMVs (**K**) of CD8α-eGFP and CD8α-mCherry; **N**, Auto- and cross-correlation functions of CD8α-eGFP and CD8α-mCherry in GPMVs; Construct scheme (**C**), Coommassie-stained SDS PAGE of recombinant His_6_-eGFP and His6-mCherry (**F**), DGS-NTA(Ni)-spiked GUV scheme (**I**) and fluorescence images of DGS-NTA(Ni) spiked GUVs with bound His_6_-eGFP and His_6_-mCherry (**L**); **O**, Auto- and cross-correlation functions of His_6_-eGFP and His_6_-mCherry bound to DGS-NTA(Ni)-spiked GUVs.

### Investigating weakly dimerizing polytopic membrane proteins by FRET

From a biophysical point of view, quantifying weaker protein-protein interactions by fluorescence techniques requires working at concentrations that are close to the dissociation constant, which may exceed levels used in single-molecule experiments. Therefore, spectroscopic techniques probing the vicinity of fluorescent probes are required, such as FRET. Working at high protein densities, with membrane-embedded entities of non-negligible excluded volume that are potentially forming dimers of unknown spatial orientation however brings specific problems, such as proximity-induced FRET, and the need to know the surface density of donors and acceptors at the site of measurement. To accommodate all the specifics of dealing with weakly interacting polytopic transmembrane proteins fused to fluorescent protein reporters, we make several basic physical and methodological considerations that will be discussed below.

### Determination of protein surface densities in the membrane of giant liposomes

Quantitative determination of dissociation constants requires simulations of FRET efficiencies by Monte Carlo approach (23), and for this, knowledge of surface densities of donors and acceptors is absolutely essential. Due to the large differences in geometry of 3D and 2D systems, calibration of fluorescence intensity versus known protein concentration feasible in 3D is not transferrable to a 2D system. Therefore, we employed a single molecule technique of fluorescence correlation spectroscopy (FCS) that enables counting fluorescent molecules within a diffraction limited spot of a radius given as 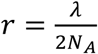 where *λ* is the excitation wavelength and *N*_A_ stands for the numerical aperture of the objective lens. We used FCS to measure fluctuations of fluorescence intensity in the upper membrane of a liposome (its apical surface) (Fig. 2A) (37), and analysis of the intensity trace yielded the absolute number of fluorophores within this focal spot. Since the FRET measurement, for which we need to know surface densities of donors and acceptors, is made in the equatorial plane of a spherical liposome (GPMV) (Fig. 2A) at a lower excitation power, the relationship between the mean equatorial intensity and the apical surface density needs to be established. FRET protein densities are much higher than the single molecule experiment requires, therefore the dependence between the equatorial signal and the apical protein density is linearly extrapolated from the FCS data (Fig. 2B).

**Fig. 2:**
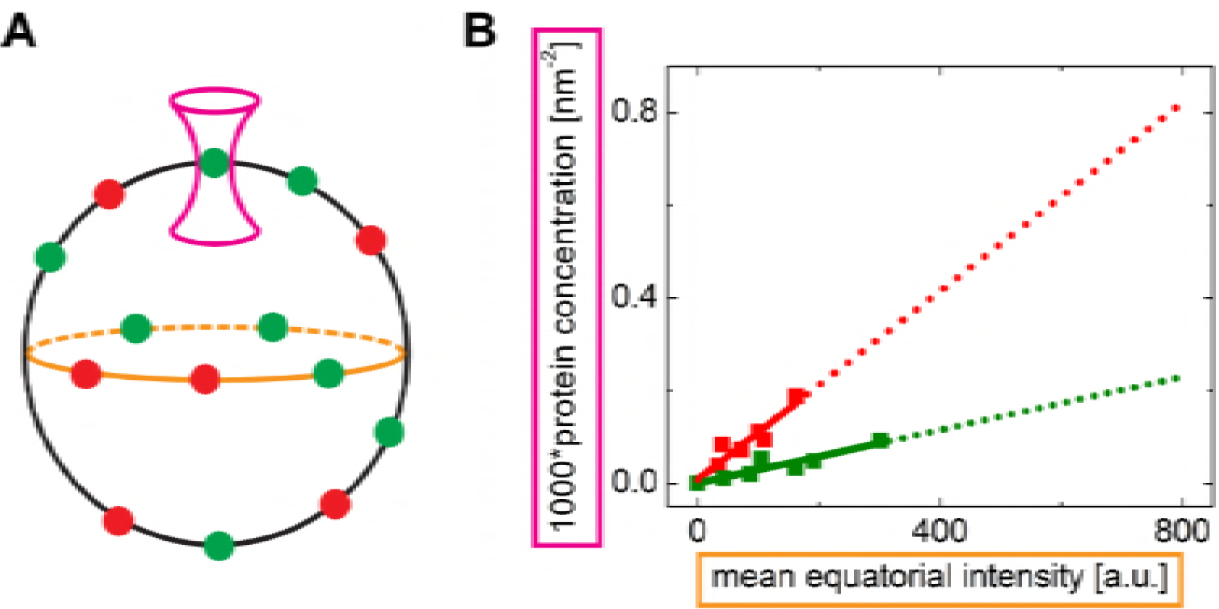
Determination of protein surface densities in the membrane of giant liposomes. (**A**) Scheme of the calibration experiment: FCCS determining the protein concentration was acquired in the apical membrane of the GUVs – the magenta laser light profile; the FRET experiment requiring the equatorial intensity of the fluorescent proteins was carried out in the middle section of the GUVs – orange ellipse. (**B**) Linear dependence of the fluorescent protein surface concentration on the mean equatorial intensity; green and red lines denote His_6_-eGFP and for His_6_-mCherry, respectively. The dotted lines show the extrapolation to the concentrations for which FCCS cannot be carried out.

### Determination of kappa squared

Förster radius represents the main characteristics of a FRET pair. It equals the distance where FRET efficiency drops to 50%, and it thus refers on the distances that can be addressed by the pair. Förster radius (*R*_0_) can be determined as follows:

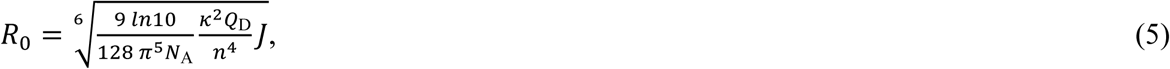

where *N*_A_ stays for the Avogadro’s number, *n* is the refractive index, *Q*_D_ is the quantum yield of the donor in the absence of acceptor, *J* is the spectral overlap integral and *κ*^2^ is the orientation factor kappa squared. This equals:

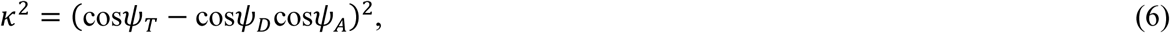

where the angles are defined by the schematics in Fig. 3A. It is typically assumed that rotational depolarization upon excitation occurs much faster than transfer of energy and that all orientations of the fluorophores are equally distributed. Such a situation is known as the isotropic dynamic limit and the value of *κ*_2_ then equals 2/3. Fluorescent proteins attached to membranes via fusions to integral membrane proteins typically rotate slowly compared to the donor fluorescence lifetime, and thus the dynamic limit no longer holds for them. In addition, membranes are naturally anisotropic, which prevents the fluorophores from sampling the entire rotational space. In this case, molecular simulations are required for the estimation of *κ*_2_ (38).

**Fig. 3:**
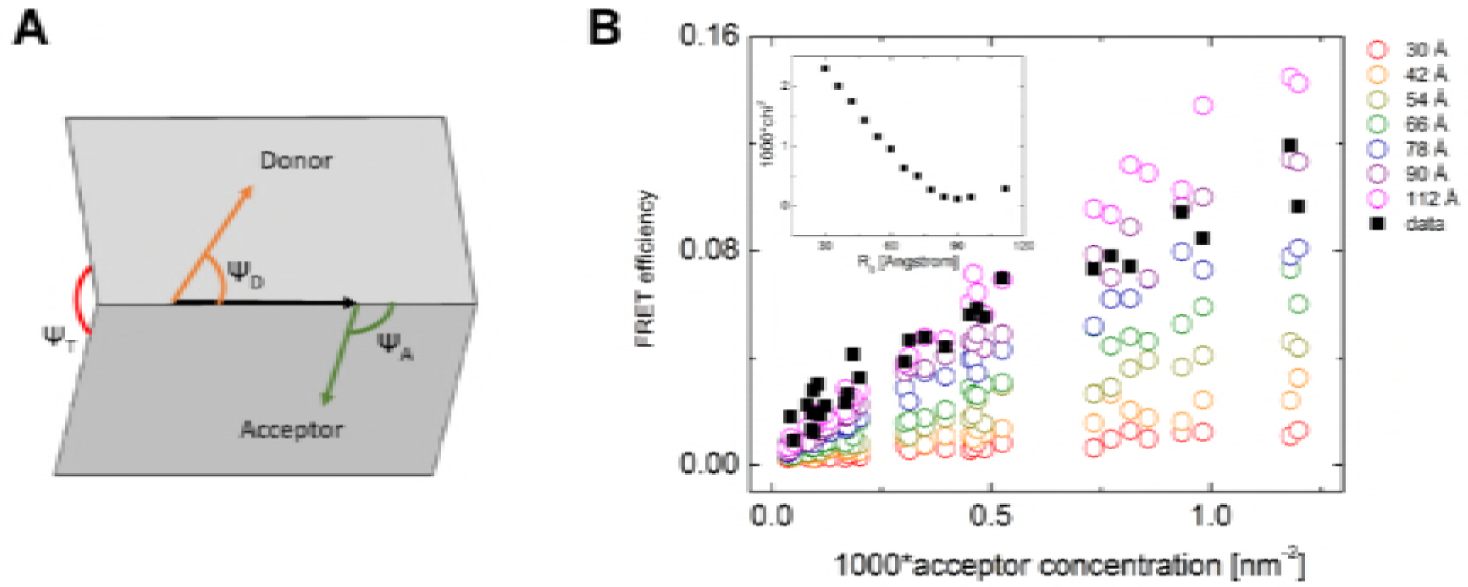
Determination of the *κ*^2^_app_ factor for two fluorescent proteins attached to the membrane. (**A**) Scheme depicting the donor/acceptor emission/absorption transition moments, and defining the angles between them. (**B**) FRET efficiency as a function of acceptor concentration: black squares denote the measured data; circles denote the simulated data with increasing apparent Förster radius. The inset shows the dependence of chi^2^ (calculated from the measured data and each of the simulated dependences) on the apparent Förster radius. The best agreement between the measured and the simulated data was found for the apparent Förster radius of 90 Å.

The *κ*^2^ factor in the static or intermediate regime depends on the donor-acceptor separation, and its average value cannot be used to calculate FRET efficiency or a FRET rate (39). Such a complex FRET kinetics would however make the quantification of FRET on membranes hardly feasible. We therefore made use of a model system that experimentally best resembles the system of our interest (GPMVs with transmembrane proteins fused to fluorescent proteins), and that is giant unilamellar vesicles (GUVs) decorated with non-interacting fluorescence proteins attached via lipid anchors. Geometrically, both systems represent planes with green and red fluorescing protein barrels attached by a short linker, i.e. the rotational motion of the barrels and their mutual geometry would be comparable. Therefore, we assume that the donor-acceptor distance-dependent *κ*^2^ factor distribution would be similar to that in GPMVs containing polytopic membrane proteins fused to fluorescence proteins, and that this distribution can be replaced by an apparent *κ*^2^_app_ factor that fulfills the following equation for FRET rate from the *i*^th^ donor:

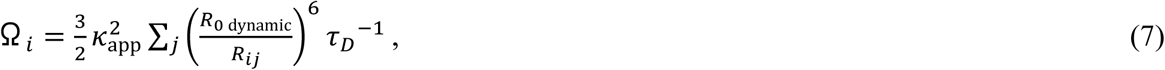

where *R*_0_ dynamic is the Förster radius that would hold for *κ*^2^ equal 2/3, and *τ*_D_ is the donor lifetime. The sum is carried out for all the acceptors present in the system. The use of this approach would be restricted only to the systems where this assumption holds, i.e. those with similar rotational dynamics and geometry. It has to be pointed out that in any FRET experiments using established interacting or non-interacting membrane proteins as positive or negative controls, this assumption is tacitly expected to hold, because otherwise the differences in FRET efficiencies could be caused by a different *κ*^2^ distribution rather than by interaction between the proteins of interest.

In order to estimate the *κ*^2^_app_ factor for two fluorescent proteins attached to the membrane, we made use of the discussed proximity FRET effect. Knowing that the distribution of His_6_-eGFP and His_6_-mCherry attached to the membrane of giant unilamellar vesicles (GUVs) spiked with DGS-NTA(Ni) is homogeneous and that no significant interaction between the two proteins occurs, the combination of FRET with MC simulations allows estimation of the apparent Förster radius of this fluorophore pair. At given surface densities of donors and acceptors, and with reasonable estimate of the excluded areas of the proteins, the apparent Förster radius is the only variable parameter required for the determination of FRET efficiency. We have thus prepared GUVs containing DGS-NTA(Ni) and measured FRET efficiency for various amounts of His-tagged fluorescent proteins added to the GUVs. For every GUV, the mean equatorial intensity was evaluated, the surface protein density of donors and acceptors was calculated from the extrapolation of the calibration measurement described above, and FRET efficiency based on fluorescence lifetime imaging (FLIM) was determined (Fig. 3B).

The surface densities of donors and acceptors obtained for each GUV were taken as input parameters for MC simulations that were employed to calculate a theoretical level of FRET efficiency for the given situation including the proximity FRET phenomenon (Fig. 3B). Varying the value of the Förster radius (*R*_0_) and assuming that the donor (eGFP) and acceptor (mCherry) cannot come closer to each other than 30-35 Å (4), we have obtained the best agreement between our experimental data and the MC simulation for *R*_0 app_ = 90 Å 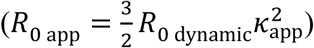 corresponding to the value of *κ*^2^_app_ ≈ 1.1, which significantly differs from the commonly used 2/3 (inset of Fig. 3B). A higher value of *κ*^2^_app_ slows down FRET rate and thus makes FRET more sensitive for longer donor-acceptor distances.

### Dimerization of RHBDL2 and CD8α

Kappa squared determination for the used donor-acceptor pair revealed that in the case of fluorescent proteins attached to the plane of the membrane, FRET enables observation of distances that are larger than those predicted by the dynamic isotropic limit. This can be advantageous if the studied protein pair is large, as FRET can in this case address also larger distances that would be invisible if the donor and acceptor were fast sampling the full rotational space. On the other hand, the proximity FRET effect becomes more pronounced, since non-interacting acceptors appear in the FRET range of the donors more frequently. Therefore, to reach an objective conclusion on the protein-protein interaction or lack thereof, simulating the theoretical FRET efficiencies at experimental protein densities is required.

To address the thermodynamic propensity of the human rhomboid intramembrane protease RHBDL2 to form dimers in the natural lipid environment, we have measured FRET in giant plasma membrane-derived vesicles (GPMVs) prepared from HeLa cells co-expressing eGFP and mCherry fusions to RHBDL2, using CD8α as a dimeric positive control (Fig. 1). The use of the spherical GPMVs was crucial for quantification of protein surface densities in the area from which the signal was collected. In living cells, membrane proteins are synthetized at the endoplasmic reticulum (ER) membranes and are then trafficked to the plasma membrane. Since plasma membrane surface is not simply planar, but it is complicated by ruffles and numerous filopodia-like protrusions (Fig. 1G, H), plenty of signal intensity heterogeneities are visible when focusing on the plasma membrane adhering to the glass (Fig. 1G, H). This could be also due to numerous ER-plasma membrane contact sites. As a consequence, overall area of the plasma membrane cannot be easily measured in live cells. Thus, GPMVs with low fluorescence signal were used for the calibration of surface protein density by FCS as described above, and protein density in the apical surface of the highly fluorescent GPMVs used for FRET measurements was calculated from the mean equatorial fluorescence. These surface protein densities were used as input parameters for MC simulations, where the value of *K*_D_ and the closest donor-acceptor distance (excluded radius) were varied, and the results were compared to the experimental values of FRET efficiency (Fig. 4).

**Fig. 4:**
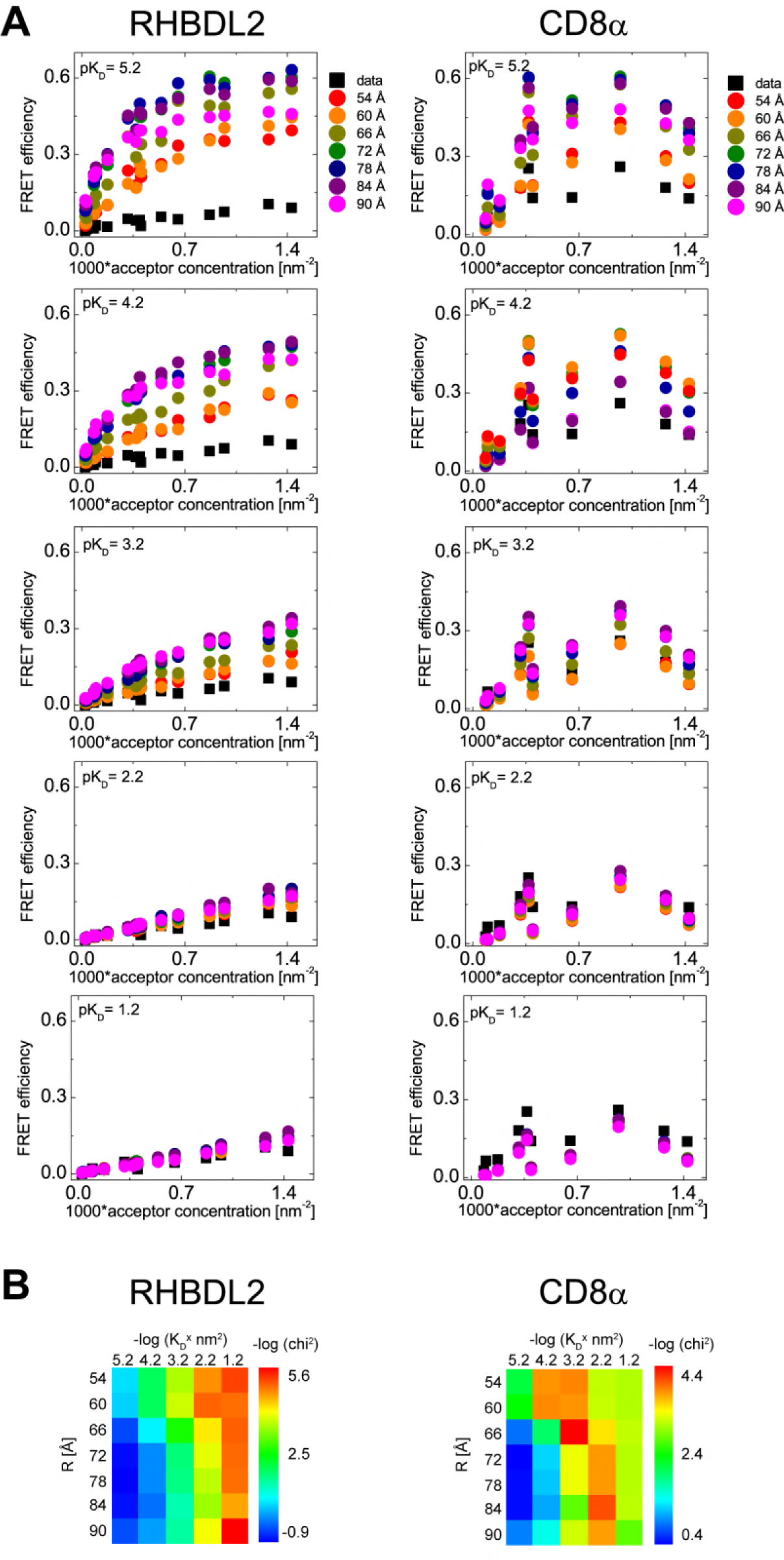
Analysis of dimerization of RHBDL2 and CD8α in GPMVs derived from HeLa cells by FRET measurements and MC simulations. (**A**) Comparison of acceptor concentration dependent FRET efficiencies obtained experimentally (black squares) and from MC simulations (circles). The MC simulations were carried out for RHBDL2 and CD8α, at increasing *K*_D_ values, and for excluded radii ranging from 54 to 90 Å. Donor concentrations are not depicted for the sake of simplicity, but for every analyzed GPMV, they were used as an input parameters for the MC simulations. (**B**) Heat-maps of −log (chi^2^) visualizing the agreement between the measured and the simulated data for various values of *K*_D_ and the excluded radii.

The results for RHBDL2 show that independently of excluded radius of the protein (R in Fig. 4B), experimental data match the simulations only at relatively high *K*_D_ values (low −log(KD×nm2)) of about 6×10−2 nm−2 (corresponding to about 2 mol % assuming lipid head group size is 60 Å^2^). This is one or two orders of magnitude higher than the highest acceptor concentration achieved in our experiment, and a few more orders of magnitude higher than the concentration of the endogenous RHBDL2 in HeLa cells (Fig. S1). In summary, it is unlikely that the rhomboid protease scaffold of RHBDL2 is intrinsically dimeric in lipid membranes at physiological concentrations. Dimerization could occur only if there was an additional force that drives the partners to one another, such as interaction via a third partner, segregation to some specific lipid pools, or binding to some cytosolic juxtamembrane structures associated with the actin cortex, but there is currently no evidence for any of these mechanisms.

For the positive control, CD8α, the simulation predicts apparent *K*_D_ value of 6×10^−4^ – 6×10^−3^ nm^−2^ (corresponding to 0.02 – 0.2 mol %, respectively, provided the lipid head group size is 60 Å^2^), which is 1-2 orders of magnitude lower than that of RHBDL2, and in agreement with the known dimeric character of this protein (36). The FRET data also explain the relatively low cross-correlation amplitude obtained from FCCS (Fig. 1), which implies the presence of CD8α dimers, but is considerably below the value expectable for a 100% dimeric protein. Together, these data indicate that some monomeric CD8α is present in GPMVs. This is consistent with the possibility that not all CD8α dimers may be disulphide-bonded in these redox conditions, and these non-covalent dimers are in dynamic equilibrium with monomers, which may be stable in the absence of endocytosis and membrane protein quality control in GPMVs.

Our analysis indicates that RHBDL2 is not dimeric to any significant degree in cell membrane derived vesicles at expression levels far exceeding the endogenous ones (17 to 56-fold higher, Fig. S1). We however could not exclude that in other intracellular compartments such as the ER or Golgi, which have a different lipid composition and hydrophobic thickness, RHBDL2 may dimerize. Since interactions within the intracellular compartments of the secretory pathway are not easily directly addressable by FRET or FCCS, we resorted to a cell biological approach exploiting the cellular mechanism of retrieving proteins back to the ER from early Golgi via the KDEL tag (40) and membrane-bound KDEL receptor (41, 42). In fact, KDEL tagging was shown to exert a strong dominant negative effect on the secretion of dimeric proteins such as TGFβ (43), documenting feasibility of this approach. When eGFP-RHBDL2 and mCherry-RHBDL2 are co-expressed in HeLa cells, both constructs show predominantly plasma membrane localization, notably labelling filamentous extrusions of the cell surface (Fig. 5A). When both of these constructs are equipped with a C-terminal, luminal KDEL tag, both show predominantly an ER localization with a complete overlap and complete loss of filopodia-like labelling (Fig. 5B). When, however, mCherry-RHBDL2-KDEL and eGFP-RHBDL2 are co-expressed, only mCherry-RHBDL2-KDEL relocalizes to the ER, while the localization of eGFP-RHBDL2 is barely influenced and fluorescence of the two reporters overlaps only minimally (Fig. 5C). The same is true when eGFP-RHBDL2 is KDEL-tagged and mCherry-RHBDL2 is not (data not shown). We therefore conclude that using this approach we do not detect formation of stable dimers of RHBDL2 that could traffic together. In other words, RHBDL2 is prevalently monomeric in all major cellular membrane compartments where it normally resides.

**Fig. 5:**
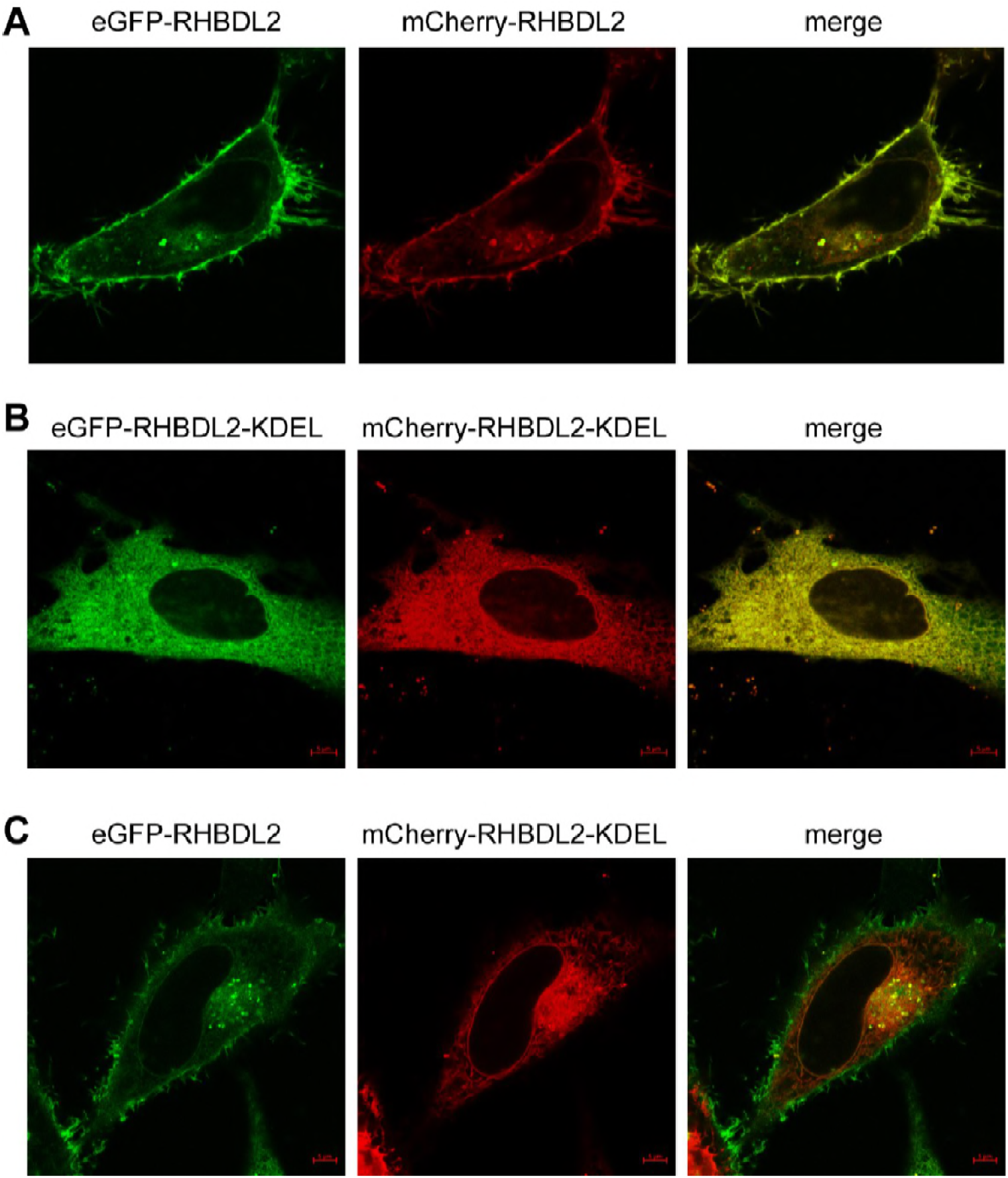
Relocalization analysis of RHBDL2 dimerization in live cells. Fluorescent constructs of human RHBDL2 fused to either eGFP or mCherry were co-expressed in HeLa cells with or without the ER-retaining KDEL signal fused to the very C-terminus of each protein, and live cell fluorescence was recorded 20-24 hrs after transfection. **A**, eGFP-RHBDL2 co-expressed with mCherry-RHBDL2; **B**, eGFP-RHBDL2-KDEL co-expressed with mCherry-RHBDL2-KDEL; **C**, eGFP-RHBDL2 co-expressed with mCherry-RHBDL2-KDEL. Note that while both fusions show strong plasma membrane localization including filopodia (A), KDEL tagging effectively relocalizes both to the ER (B), while KDEL tagging of only one of them does not relocalize the other fusion (C), meaning that the two fusion proteins do not stably interact with one another within the cell.

## Discussion

For addressing weak protein oligomerization in cellular membranes, FRET combined with FLIM represents one of a few applicable optical non-invasive techniques. It has been shown that the expression levels of fluorescent protein acceptor fusion has to be kept under control as it may contribute to the overall FRET efficiency by the proximity effect (2). Having full control of lateral protein density does not seem to be possible in living cells, but it is possible in GPMVs derived from the cells of interest using an external calibration (4). To separate the impact of the proximity effect from the impact caused by protein oligomerization on the overall FRET efficiency we have utilized MC simulations.

A large fluorescent protein attached to a membrane-residing protein of interest, however, i) cannot freely rotate in all directions, and ii) rotates on the time-scale much slower that the donor fluorescence lifetime. Overall, these fluorescent fusion membrane proteins do not fulfil the requirements for the dynamic isotropic limit which predicts the value of the orientation factor *κ*^2^ to be 2/3. Knowledge of *κ*^2^ is however essential for meaningful quantification of FRET. In situations that are neither isotropic nor dynamic, estimation of *κ*^2^ is very difficult and usually requires understanding the dynamics of the involved proteins. Moreover, to calculate FRET efficiency under the static or intermediate conditions, the overall dependence of the *κ*^2^ factor on the donor-acceptor distance has to be known (39). Here, we propose to use an apparent *κ*^2^_app_ value that would allow to compare systems resembling each other well as regards rotational dynamics and geometry of the fluorophores. In this sense, we consider our system consisting of transmembrane proteins fused to fluorescence proteins comparable to the eGFP/mCherry donor/acceptor pair attached by His-tag to the membrane of GUVs. Our approach then simulates FRET efficiencies at experimentally obtained protein densities and compares those with the measured FRET efficiency values. The only optimized parameter in the simulations is the apparent Förster radius that comprises the value of *κ*^2^.

Our results clearly indicate that the apparent Förster radius for eGFP and mCherry in our system is around 90 Å, which is significantly larger than commonly reported in literature, i.e. 52 Å (44). This makes FRET ‘see’ much longer distances in 2D than would be expected. Combination of this fact with the 2D membrane confinement significantly complicates drawing conclusions on oligomerization based solely on FRET observation as this would occur purely as a result of protein density, and the use of MC simulations is needed for interpretation of the FRET data.

Using this toolset, we addressed the dimerization propensity of human intramembrane protease RHBDL2. The FRET efficiencies obtained in the membranes of GPMVs derived from cellular membranes were compared with the efficiencies obtained from the MC simulations, which revealed no protein-protein interaction at the density level we studied, which is far above the endogenous one. In order to validate our method, we have also applied it to the dimeric membrane protein CD8α. The data indicated that CD8α was dimeric in GPMVs, and the values of the dissociation constant of CD8α were 1-2 orders of magnitude lower than those of RHBDL2. Interestingly, the *K*_D_ value of CD8α is very similar to the *K*_D_ value reported for glycophorin A (4), a model dimeric transmembrane proteins. Strikingly, both CD8α and glycophorin A are commonly thought to be dimeric, but our data on CD8α and the data of others glycophorin A (4) indicate that a significant portion of the population of both proteins are monomeric in cell-derived GPMV membranes. In the case of CD8α, which is a disulphide-linked dimer in cells, it is possible that the redox environment in and around GPMVs leads to the partial reduction of the disulphides and partial monomerization. The discrepancy between the expected aggregation state of glycophorin A known from previous reports and measured by FRET in GPMVs has been related to the complexity of the cellular membrane and protein crowding that may limit the dimerization propensity of glycophorin A (4), which could apply also to CD8α in addition.

Furthermore, our simulations show an interesting and unintuitive behavior of the dependence of maximum FRET efficiencies (at saturating acceptor concentration) on the donor-acceptor separation (see the differently colored series of data-points in the upper left panel of Fig. 4A for an example). From the lowest simulated values of donor-acceptor separation (54 Å in our case) upwards, the maximum FRET efficiency at saturating acceptor concentration increases with the separation until it reaches a maximum at ~78 Å and then starts decaying. This is counterintuitive, because it is generally expected that the farther the acceptor is from the donor (i.e. the higher the separation), the lower FRET efficiency is. However, in our system with the apparent Förster radius increased to about 90 Å, we deal with donor-acceptor separations that are much below this value. At this distance regime, FRET efficiency decreases with distance only moderately even for a single donor/acceptor pair. The phenomenon we observe in our MC simulations implies, more generally, that in a system with high confinement and large proximity contribution, such as ours, a distance regime can be found where the influence imposed by the overall arrangement of the acceptors in space prevails over the anticipated simple distance dependence. As a result of that, quantification of the experiment including the determination of *K*_D_ and the excluded radius inevitably requires Monte Carlo simulations, which take into account all possible kinds of energy transfer, not only between the donors and the acceptors within a dimer, between free monomers, but also between the donors and acceptors from distant dimers. The obvious implication of our findings is that for transmembrane proteins in lipid membranes, a simple comparison of FRET data with control interacting/non-interacting protein pairs of different geometry is insufficient to draw conclusions on protein-protein interactions.

## Conclusions

Here we adapt the usage of FRET for the analysis of dimerization of polytopic α-helical transmembrane proteins by taking into account proximity induced FRET, by estimating the kappa squared parameter that significantly deviates from the limiting value of 2/3 commonly used in isotropic conditions, and by employing Monte Carlo simulation to interpret FRET results. Using these methods, our biophysical and cell-biological experiments do not provide any evidence for the dimerization of human rhomboid protease RHBDL2 in lipid membranes derived from live cells. This suggests that in lipid membranes, the transmembrane core of rhomboid protease may be intrinsically monomeric, unlike proposed previously (13). If this is indeed the case more widely, the proposed allosteric activation of rhomboids by their dimerization reported previously (13) is not a native general property of these enzymes, and it could be simply a detergent artifact. Specifically for the understanding of the mechanisms and biology of rhomboid superfamily members, it is important to scrutinize whether other rhomboid proteases or other members of the rhomboid superfamily, such as iRhoms and Derlins can dimerize in lipid membranes at physiologically relevant concentrations, or whether they are also intrinsically monomeric.

## Author Contributions

KS, JH, and JŠ designed research, JŠ, JH, PR, LA, AS, DJ and EP performed research, JH, JŠ, DJ and KS analysed data, and JH and KS wrote the manuscript.

## Acknowledgements

We thank Evzen Bouřa for the His_6_-eGFP and His_6_-mCherry encoding plasmids, and Marek Cebecauer and Radek Šachl for critical reading of the manuscript. KS acknowledges support from Czech Science Foundation (project no. 18-09556S), Ministry of Education, Youth and Sports of the Czech Republic (project no. LO1302) and European Regional Development Fund; OP RDE (No. CZ.02.1.01/0.0/0.0/16_019/0000729). Authors state no conflicts of interest with the content of this article.

## Figures with Legends

**Fig. S1:**
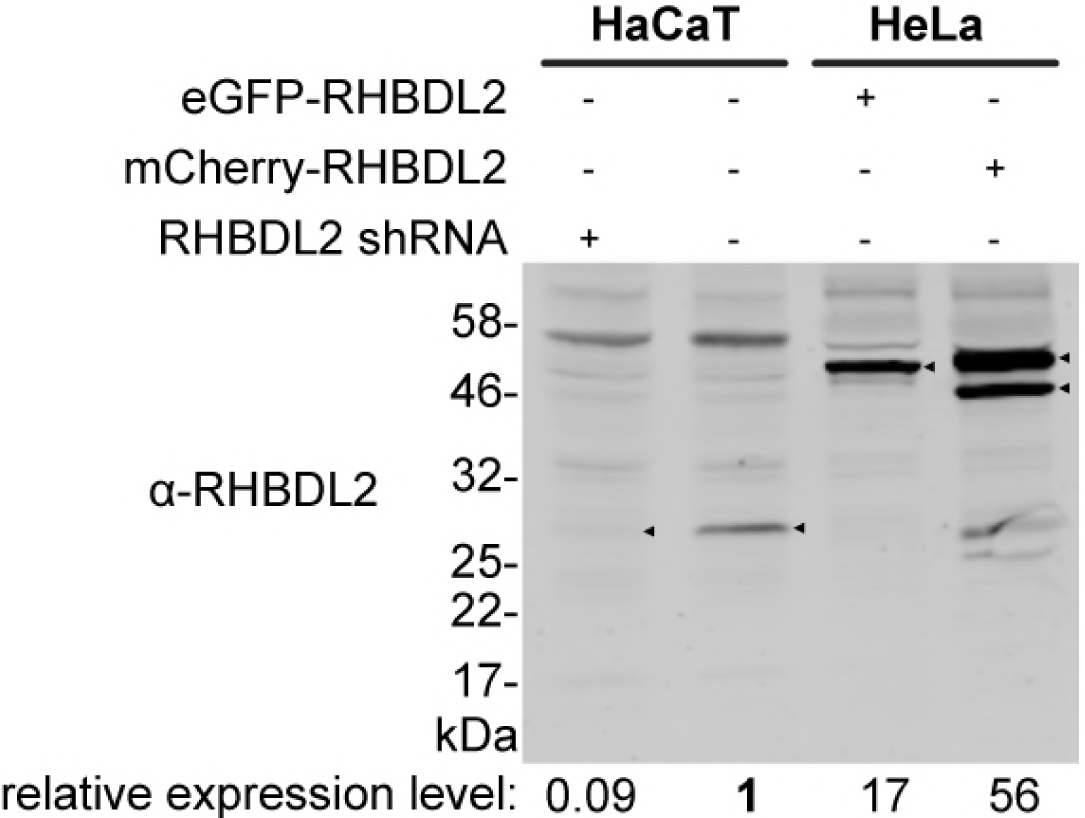
Comparison of expression levels of transfected eGFP/mCherry-RHBDL2 to the levels of endogenous RHBDL2. Approximately equal amounts of cell lysates of HeLa cells transiently transfected by eGFP/mCherry-RHBDL2 and human keratinocytes HaCaT with or without expressed shRNA targeting endogenous RHBDL2 were separated by SDS PAGE and analyzed by quantitative immunoblotting using α-RHBDL2 primary antibody and fluorescent secondary antibody as described in Methods. Fluorescence of the secondary antibody was visualized using infrared scanner, and expression levels of RHBDL2 were quantified from the integrated fluorescence intensity values summed up for the specific bands (marked by black triangles) and normalized for the total protein in each lane indicated by the fluorescence intensity of the Revert staining (Methods).

